# DAPK1-mediated parkin inactivation enhances neurotoxicity via MITOL-dependent degradation

**DOI:** 10.64898/2026.01.07.698295

**Authors:** Chul Hong Park, Donghyuk Shin, Kwang Chul Chung

**Author notes:** To whom correspondence should be addressed: Department of Systems Biology, College of Life Science and Biotechnology, Yonsei University, Yonsei-ro 50, Seodaemun-gu, Seoul 03722, Korea.

## Abstract

Parkinson’s disease (PD) is characterized by progressive neurodegeneration and is marked by the formation of Lewy bodies, which are intracellular aggregates primarily composed of α-synuclein. Mitochondrial dysfunction and impaired protein degradation pathways play critical roles in the progression of PD, contributing to the loss of dopaminergic neurons in the *substantia nigra*. Phosphorylation of α-synuclein promotes its aggregation, underscoring its role in disease progression. Parkin, an E3 ubiquitin ligase, is considered to be a pleiotropic, neuroprotective protein that modulates the mitochondrial quality control as well as metabolic turnover and the accumulation of α-synuclein. Death-associated protein kinase 1 (DAPK1), involved in controlling apoptosis and autophagy, has recently emerged as an important factor in neurodegeneration. While DAPK1 is implicated in Alzheimer’s disease through its role in tau aggregation and amyloid-β production, we demonstrate that DAPK1 plays a role in PD by phosphorylating parkin at Ser136 and Ser198. This phosphorylation causes the mitochondrial transport of parkin, enhancing interaction with mitochondria-localized E3 ubiquitin ligase MITOL and consequently leading to the degradation of parkin. As parkin is critical for neuroprotection, its degradation exacerbates the toxic effect of 6-hydroxydopamine, further compromising neuronal survival. These results indicate that DAPK1 acts as a previously unrecognized modulator of parkin and a key contributor to PD pathogenesis, bridging pathways of mitochondrial dysfunction, α-synuclein aggregation, and neuronal cell death.

## Introduction

Parkinson’s disease (PD) is a progressive neurodegenerative disorder characterized by the selective degeneration of dopaminergic neurons in the substantia nigra and the accumulation of intracellular inclusions known as Lewy bodies (LBs) (Spillantini et al., 1997). The primary component of LBs is α-synuclein, a presynaptic protein that is normally soluble but undergoes misfolding and aggregation in PD (Wakabayashi et al., 2007). The pathological aggregation of α-synuclein is closely linked to its phosphorylation, primarily at Ser129 (Fujiwara et al., 2002; Anderson et al., 2006). This phosphorylation event promotes α-synuclein fibrillation and is considered a hallmark of PD pathology (Tenreiro et al., 2014). In addition to α-synuclein aggregation, mitochondrial dysfunction and impaired protein degradation pathways further exacerbate PD pathogenesis by increasing oxidative stress, promoting neuroinflammation, and disrupting cellular homeostasis (Schapira et al., 2011; Youle et al., 2011). Defects in mitochondrial quality control mechanisms, such as mitophagy, have been strongly implicated in PD progression, particularly through mutations in key genes such as PINK1 and parkin (Narendra et al., 2008; Pickrell & Youle, 2015).

Parkin is an ubiquitin (Ub) E3 ligase, which promotes the degradation of target proteins through polyubiquitination or modulates their biochemical properties via mono-ubiquitination. Therefore, parkin plays a critical role in neuronal survival by regulating protein homeostasis and cellular stress responses (Koyano et al., 2014). Various forms of parkin mutation lead to early-onset PD, highlighting its neuroprotective role (Henrich et al., 2023). Like many other proteins playing crucial roles of cellular functions and regulation, parkin is also subjected to various forms of post-translational modification (PTM), including phosphorylation, ubiquitination, covalent modification of ubiquitin-like modifiers. For example, dual specificity protein kinase, Dyrk1A, phosphorylated parkin at Ser131, causing the inhibition of its E3 ubiquitin ligase and neuroprotective activity (Im & Chung, 2015). In addition, covalent modification with NEDD8, an ubiquitin-like posttranslational modifier, positively regulates the E3 ligase activity of parkin, potentiating its cytoprotective function against 1-methyl-4-phenylpyridinium ion in dopaminergic neuronal cells (Um et al., 2012).

Death-associated protein kinase 1 (DAPK1) is a calcium/calmodulin-dependent Ser/Thr kinase involved in various cellular processes, including apoptosis, autophagy, and cytoskeletal regulation (Bialik & Kimchi, 2006). To date, many substrates of DAPK1 have been identified, and its activity has been shown to influence a wide range of cellular functions by modulating the biochemical properties and functional activities of these substrates. Notably, DAPK1 regulates toxic protein aggregation in the brain, and alterations in its kinase activity are closely associated with neurodegenerative diseases (NDDs) (Wang et al., 2016). For example, DAPK1 has been extensively studied in the context of Alzheimer’s disease (AD), where it promotes tau phosphorylation and amyloid-β accumulation, contributing to neuronal toxicity (Duan et al., 2013). These findings further suggest that DAPK1 may also play a role in the onset and progression of PD, a hypothesis supported by several recent reports, including those from our lab. In PD, DAPK1 expression is significantly upregulated in both human post-mortem PD brains and animal models, and this upregulation correlates with increased α-synuclein phosphorylation and dopaminergic neuronal loss (Su et al., 2019). We have also demonstrated that DAPK1 directly phosphorylates α-synuclein at Ser129, enhancing its aggregation and toxicity (Shin & Chung, 2020).

The previous findings, which highlight a molecular interaction and functional relationship between parkin and α-synuclein, both of which are closely involved in the formation of LBs in PD, have led us to hypothesize a functional relationship between DAPK1 and parkin. In this study, we sought to explore this hypothesis further. Beyond its role in α-synuclein pathology, our study provides new evidence that DAPK1 also targets parkin. We further demonstrate that DAPK1 phosphorylates parkin at Ser136 and Ser198, impacting its stability and function. This phosphorylation event increases neuronal vulnerability to oxidative stress, contributing to PD-related neurodegeneration. These findings suggest that DAPK1 contributes to PD pathogenesis not only by promoting α-synuclein aggregation but also by enhancing neuronal susceptibility to oxidative stress through its regulation of parkin. In summary, our study identifies DAPK1 as a key regulator of PD pathogenesis through its dual role in α-synuclein pathology and neuronal vulnerability.

## Materials and Methods

### DNA constructs and RNA interference

FLAG-tagged mammalian expression plasmids carrying the cDNA of human DAPK1 in its wild-type form (FLAG-DAPK1-WT) or a kinase-deficient mutant (FLAG-DAPK1-K42A, also termed FLAG-DAPK1-KD) were generously provided by T.H. Lee (Fujian Medical University, Fujian, China). A pcDNA3.1 construct expressing Myc-tagged human parkin (pcDNA3.1-Myc-Parkin) was kindly obtained from K. Tanaka (Tokyo Metropolitan Institute of Medical Science, Tokyo, Japan). Bacterial vectors encoding GST-fused N-terminal parkin, either wild type or carrying point mutations (S131A, S136A, or S198A), were generated as previously described (Im & Chung, 2015). The double-mutant version (S136A/S198A) was generated by site-directed mutagenesis of pcDNA3.1-Myc-Parkin. Truncated fragments of parkin (Myc-parkin-1–225 and Myc-parkin-226–465) were amplified with PrimeSTAR HS DNA polymerase (TAKARA, Shiga, Japan) and subsequently inserted into the pRK5-Myc vector. Constructs expressing HA-tagged ubiquitin mutants, in which either lysine 48 (pRK-HA-Ub-K48) or lysine 63 (pRK-HA-Ub-K63) was preserved while all remaining lysine residues were substituted with arginine, were obtained from Addgene (Cambridge, MA, USA). FLAG-tagged human MITOL expression plasmids, including both the wild-type protein (MITOL-WT) and a catalytically inactive form (MITOL-MT), were provided by S. Hirose (Tokyo Institute of Technology, Tokyo, Japan). siRNAs specific for DAPK1 and a scrambled negative control were obtained from IDT Korea (Hanam-si, Gyeonggi-do, Korea). The sequences of the *DAPK1*-specific siRNA duplex were as follows: sense, 5′-GGUGAGGCGUGACAGUUUAUCAUGA(dTdT)-3′; antisense, 5′-AACCACUCCGCACUGUCAAAUAGUACU(dTdT)-3′.

### Cell culture and DNA transfection

Human embryonic kidney 293 (HEK293) cells were grown in Dulbecco’s Modified Eagle Medium (DMEM) supplemented with 10% fetal bovine serum (FBS) and 100 U/ml penicillin–streptomycin. Mouse embryonic fibroblasts (MEFs) obtained from either wild-type *(DAPK1*+/+) or knockout (*DAPK1*−/−) mice were generously provided by T.H. Lee (Fujian Medical University, Fujian, China) and cultured in Dulbecco’s Medium containing 10% FBS and antibiotics (1% penicillin - streptomycin). Mouse neuroblastoma MN9D cells were maintained in high-glucose DMEM with 10% FBS and 1% penicillin - streptomycin on dishes pre-coated with poly-D-lysine (25 μg/ml; Sigma-Aldrich), under atmospheric conditions of 90% air and 10% CO₂. Unless specified otherwise, all cell lines were propagated at 37°C in a humidified incubator with 5% CO₂. Transient transfections were performed using either Lipofectamine 2000 or polyethyleneimine (PEI) following the manufacturers’ standard instructions.

### Cell line information and authentication

HEK293 cells (*Homo sapiens*, female, embryonic kidney; RRID: CVCL_0045), MN9D cells (*Mus musculus*, male, embryonic midbrain dopaminergic neurons; RRID: CVCL_M067), and wild-type (*DAPK1*+/+) or DAPK1-deficient (*DAPK1*−/−) mouse embryonic fibroblasts (MEFs, *Mus musculus*, embryonic fibroblasts) were used in this study. MEFs were generated in-house from mouse embryos and therefore do not have an assigned RRID. All cell lines were obtained either from accredited repositories, such as ATCC and KCLB, or were kindly provided by collaborating laboratories acknowledged in this manuscript. Although in-house authentication using STR or SNP profiling was not performed, all cell lines were obtained from institutions authorized to conduct such validations. According to the Cellosaurus and ICLAC databases, none of the cell lines used in this study have been reported as misidentified or cross-contaminated. Routine mycoplasma testing was conducted using the e-MYCO™ Mycoplasma Detection Kit ver. 2.0 (iNtRON Biotechnology, Seongnam-si, Gyeonggi-do, Korea), and all cell lines consistently tested negative throughout the experimental period. All experiments were conducted between October 2022 and June 2025, using cells at passages between 5 and 10.

### Co-immunoprecipitation (Co-IP) and immunoblot analysis

Cells were washed by phosphate-buffered saline, collected by scraping, and lysed in buffer (50 mM Tris, 10% glycerol, 1% Nonidet P-40, 150 mM NaCl, 1 mM EGTA, 1 mM sodium orthovanadate, 1 mM sodium fluoride, 0.2 mM phenylmethylsulfonyl fluoride, and 1 µg/ml aprotinin). The lysates were sonicated and then cleared by centrifugation at 13,000 × g for 15 min at 4°C. For immunoprecipitation, 500–1000 µg of protein was mixed with 1 µg of the appropriate antibody and incubated overnight at 4°C with constant rotation. Protein-A Sepharose beads were subsequently added and allowed to bind for 2 h at 4°C. After several washes with lysis buffer, the immune complexes were released by heating in 2× SDS sample buffer. Samples were then subjected to SDS-PAGE, and proteins were transferred onto nitrocellulose membranes (Whatman, GE Healthcare Life Sciences). Membranes were first incubated at room temperature for 1 h in TBST buffer containing 0.1% Tween 20, 137 mM NaCl, and 20 mM Tris (pH 7.5) supplemented with 5% nonfat milk to block nonspecific binding. After blocking, membranes were incubated overnight at 4°C with the designated primary antibody. After the membranes were rinsed, they were incubated with horseradish peroxidase–conjugated secondary antibodies for 2 hours, and protein expression was visualized via enhanced chemiluminescence, as recommended by the manufacturer.

### Immunostaining analysis

MN9D neuroblastoma cells were plated onto coverslips pre-treated with poly-D-lysine. Following two rinses with PBS, the cells were incubated in 3.7% formaldehyde at room temperature for 10 min, then permeabilized using 0.1% Triton X-100 for an additional 10 min Blocking was then performed for 1 h in TBST containing 1% BSA. The cells were then incubated with rabbit polyclonal anti-FLAG and/or mouse monoclonal anti-Myc antibodies, followed by Alexa Fluor 488–or 594–labeled secondary antibodies. Fluorescent microscopy images were collected on a Zeiss LSM 880 system (Carl Zeiss, Germany) and analyzed afterward using the LSM Image Browser tool.

### Preparation of mouse whole-brain lysates

Brain tissues were isolated from 6-week-old male C57BL/6 mice (Orient Bio, Seongnam-si, Gyeonggi-do, Korea) and disrupted in lysis buffer (150 mM NaCl, 0.5% sodium deoxycholate, 50 mM Tris-HCl, 0.1% SDS, 1% Triton X-100, and a protease inhibitor). The homogenates were sonicated and then centrifuged at 4°C for 20 min at 13,000 ×g, after which the clarified supernatants were obtained and used for downstream analyses.

### Fractionation of Cellular Proteins into Triton X-100 -Soluble and -Insoluble Pools

Cells were rinsed twice with ice-cold PBS and lysed in a buffer containing 150 mM NaCl, 10 mM Tris-HCl (pH 7.4), 1% Triton X-100, 10% glycerol, 20 mM N-ethylmaleimide, and protease inhibitors. The lysates were centrifuged at 4°C, 15,000 × g for 20 minutes, and the supernatants were collected as the Triton X-100–soluble fraction. The resulting pellets were rinsed, suspended in lysis buffer with 4% SDS and incubated at elevated temperature for 30 min to solubilize proteins. Both fractions were subjected to SDS-PAGE and subsequently analyzed by immunoblotting.

### Purification of recombinant GST–parkin from bacteria

*E. coli* BL21 cells carrying GST–parkin plasmids were grown until the culture reached an OD_600_ of approximately 0.7–0.8. Protein expression was initiated by adding 0.5 mM IPTG (Sigma-Aldrich), and cultures were maintained at 37°C for 24 h. Bacterial pellets were collected and disrupted on ice by sonication in a buffer composed of 50 mM Tris-HCl (pH 7.4), 200 mM NaCl, 1 mM EDTA, 1 mM DTT, 0.1% Triton X-100, and a protease inhibitor cocktail. The lysates were cleared by centrifugation at 12,000 × g for 20 min, and the supernatant fraction was incubated with glutathione Sepharose 4B beads (GE Healthcare Life Sciences) overnight at 4°C. After thorough washing of the beads, bound GST–parkin proteins were released using buffer containing 10 mM reduced glutathione and 50 mM Tris-HCl (pH 7.4).

### In vitro kinase assay

MN9D cells were transfected for 24 h with either FLAG-DAPK1-WT or FLAG-DAPK1-KD plasmids. Cell lysates prepared in NP-40 buffer were subjected to immunoprecipitation using anti-FLAG antibody overnight at 4°C, followed by capture with protein A–Sepharose beads for 2 h. After washing with lysis buffer, the beads were further washed using kinase buffer composed of 50 µM DTT, 20 mM MgCl₂, and 40 mM Tris-HCl (pH 7.5). The resulting immune complexes were incubated with 2 µg of recombinant GST–parkin in kinase buffer supplemented with 10 µM ATP. Kinase activity was initiated by the addition of 10 µCi [γ-^32^P] ATP, and the reactions were allowed to proceed at 30 °C for 30 min. Reactions were stopped by adding SDS sample buffer, and phosphorylation of substrates was examined by SDS-PAGE followed by autoradiography.

### Cell viability assay

Cells transfected with DNA or siRNA were treated with or without 6-hydroxydopamine (6-OHDA) for 24 h. The CCK-8 assay (Dojindo Laboratories, Kumamoto, Japan) was performed to evaluate cell viability. Culture medium was replaced with a 1:10 dilution of CCK-8 reagent in complete medium and incubated for 30 min at 37°C. Using a microplate reader, the absorbance at 450 nm was recorded.

### Alphafold prediction of Ser136-phosphorylated Parkin structure

To explore structural changes in parkin, we employed the AlphaFold3 prediction server (https://alphafoldserver.com/about). The full-length parkin sequence was submitted with serine 136 modified to phospho-serine. The server generated five highly similar structural models, and the model considered most representative was subsequently examined using PyMOL (The PyMOL Molecular Graphics System, Version 3.0, Schrödinger, LLC).

### Statistical analysis

Data were analyzed using unpaired Student’s t-tests to determine statistical significance between groups. Graphs were prepared with GraphPad Prism 5.0 (GraphPad Software, Inc.), and results are expressed as the mean ± standard deviation (SD). Densitometric quantification of Western blot bands was performed with ImageJ software (version 1.53).

## Results

### Physical interaction between DAPK1 and parkin in mammalian cells

Recent studies identified DAPK1 as a regulator of synucleinopathy and dopaminergic neuron loss in a PD mouse model (Su et al., 2019). To explore its relationship with parkin, we compared their expression individually and in combination. While DAPK1 levels remained stable, parkin levels significantly decreased upon co-expression, suggesting DAPK1 may promote parkin degradation (Fig. 1A). This reduction was reversed by proteasome inhibition with MG132 (Fig. 1B). Co-IP in HEK293 cells confirmed a specific DAPK1-parkin interaction with or without MG132 (Fig. S1 and Fig. 1B). Endogenous interactions were also detected in MN9D cells (Fig. 1C) and mouse brain lysates (Fig. 1D), with immunostaining showing colocalization of DAPK1 and parkin in MN9D cells (Fig. 1E). A GST pull-down assay using purified recombinant proteins confirmed a direct biochemical interaction (Fig. 1F). Together, these results demonstrate that DAPK1 physically interacts with parkin in mammalian cells and may regulate its stability through the proteasome pathway.

**Fig. 1.**
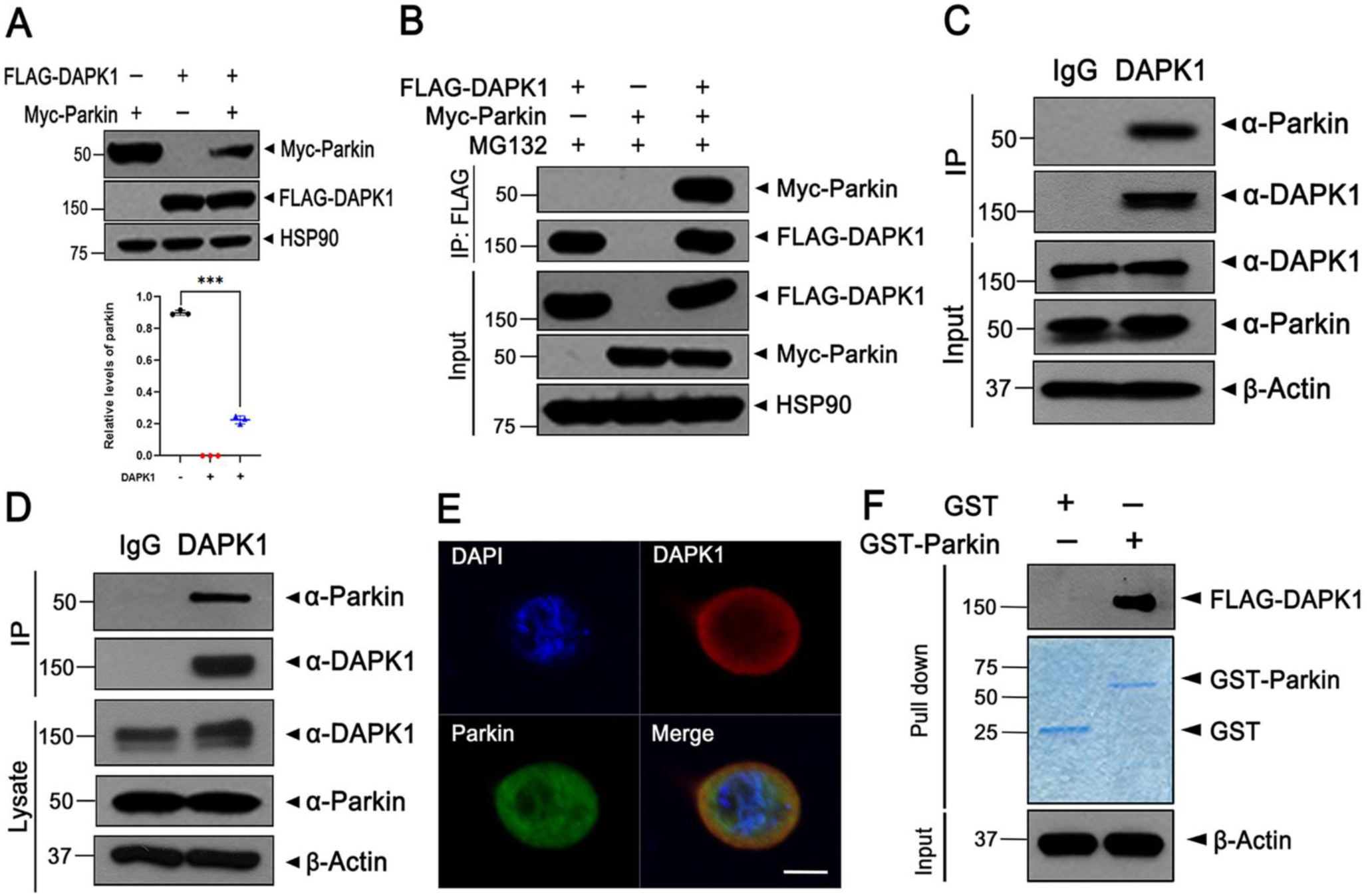
Parkin interacts with DAPK1 in mammalian cells. (A) HEK293 cells were transfected for 24 h with plasmids encoding Myc-parkin, FLAG-DAPK1, or both, as indicated. After harvest, cell lysates were immunoblotted using the indicated antibodies. Quantification of relative parkin expression was quantified (mean ± SD, n = 3; \*\*\**p* ≤ 0.0001). (B) HEK293 cells expressing Myc-parkin and/or FLAG-DAPK1 were further treated with 10 μM MG132 for 6 h before fixation. Cell lysates were subjected to immunoprecipitation using an anti-FLAG antibody, and the immunoprecipitates were analyzed by immunoblotting with the indicated antibodies. (C) Lysates prepared from MN9D cells were immunoprecipitated using either anti-DAPK1 antibody or normal IgG, and immunoblot analysis was performed using the indicated antibodies. (D) Endogenous interaction between parkin and DAPK1 was also examined in mouse brain extracts by immunoprecipitation with anti-DAPK1 or IgG, followed by immunoblotting. (E) MN9D cells were fixed, permeabilized, and subjected to immunofluorescence staining. Confocal microscopy images display colocalization of endogenous DAPK1 (red) and parkin (green), while DAPI (blue) was used to stain nuclei, and the images include scale bars of 10 μm. (F) GST pull-down assays were conducted with purified GST or GST-parkin proteins incubated with lysates from FLAG-DAPK1–expressing MN9D cells. After washing, bound proteins were detected by immunoblotting with anti-FLAG antibody. Input GST proteins were confirmed by Coomassie brilliant blue staining (blue panel). Hsp90 and β-actin served as internal controls for protein loading.

### DAPK1 promotes the degradation of parkin

We next examined the functional relationship between DAPK1 and parkin. While parkin did not affect DAPK1 levels, DAPK1 reduced parkin expression in a dose-dependent manner (Fig. S2A), suggesting DAPK1 promotes parkin degradation, possibly involving phosphorylation given its kinase function. To assess the role of DAPK1’s kinase activity, MN9D cells were transfected with Myc-parkin and either wild-type DAPK1 (DAPK1-WT) or a kinase-defective mutant (DAPK1-K42A). Parkin levels were significantly reduced by DAPK1-WT but not by DAPK1-K42A, indicating that DAPK1’s kinase activity is essential for parkin degradation (Fig. S2B). Consistent with this, endogenous parkin levels were elevated in DAPK1-knockout MEFs, while reintroduction of DAPK1 restored the reduction in parkin expression (Fig. S2C). Cycloheximide (CHX) chase assays further revealed that DAPK1-WT markedly accelerated the degradation of parkin protein compared to control, as evidenced by a significant reduction in parkin levels over time (Fig. S2D). In contrast, the kinase-inactive DAPK1-K42A mutant failed to alter parkin turnover, and the half-life of parkin remained comparable to that of the control group (Fig. S2E). These findings confirm that DAPK1 promotes parkin degradation in a kinase activity-dependent manner by reducing its protein stability

### DAPK1-mediated proteolysis of parkin occurs through ubiquitin-proteasome system

Given that eukaryotic cells degrade proteins via the ubiquitin-proteasome system (UPS) or lysosome-mediated autophagy (Ciechanover, 2005). we investigated the pathway responsible for DAPK1-mediated parkin degradation. As shown in Fig. 1B, proteasome inhibition by MG132 reversed the DAPK1-induced reduction in parkin. To confirm this, MN9D cells co-expressing DAPK1 and parkin were treated with MG132, rapamycin, or NH₄Cl. Only MG132 significantly restored parkin levels (Fig. 2A), and similar effects were observed with epoxomicin, another proteasome inhibitor (Fig. 2B), indicating UPS involvement.

**Fig. 2.**
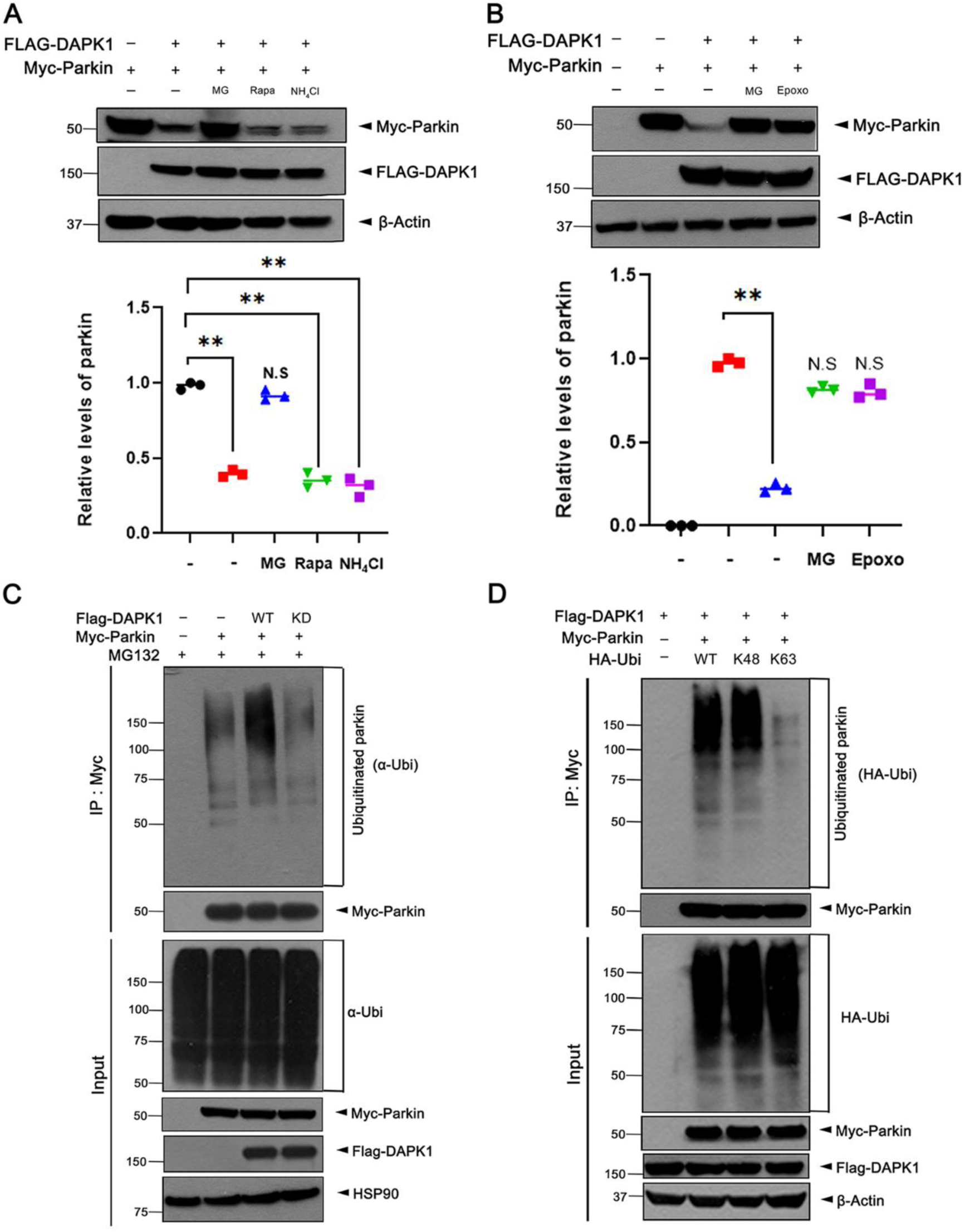
DAPK1 promotes the degradation of parkin through ubiquitin proteasome system. (A, B) MN9D cells were transfected for 24 h with Myc-parkin alone or together with FLAG-DAPK1, followed by treatment for 6 h with vehicle (–), 10 μM MG132 (MG), 1 μM rapamycin (Rapa), 25 mM NH_4_Cl, or 10 μM epoxomicin (Epoxo). Lysates were immunoblotted with the indicated antibodies. Quantification of relative parkin expression was quantified (mean ± SD, n = 3; \*\**p* ≤ 0.001; N.S., not significant). (C) MN9D cells were transfected for 24 h with FLAG-DAPK1 alone or co-transfected with Myc-parkin, then incubated with 10 μM MG132 for 6 h. Cell lysates were immunoprecipitated using anti-Myc antibody, and the precipitates were analyzed by immunoblotting. (D) Where indicated, MN9D cells were co-transfected for 24 h with FLAG-DAPK1, Myc-parkin, and either HA-ubiquitin-WT, HA-ubiquitin-K48, or HA-ubiquitin-K63. After additional incubation with 10 μM MG132 for 6 h, cell lysates were immunoprecipitated with anti-Myc antibody, and the resulting immunoprecipitates were examined by immunoblotting. Hsp90 and β-actin served as internal controls for protein loading.

We next examined whether DAPK1 promotes parkin ubiquitination. *In vitro* ubiquitination assays showed that DAPK1-WT, but not the kinase-dead DAPK1-KD, enhanced parkin ubiquitination, confirming kinase-dependent regulation (Fig. 2C). Since K48-linked polyubiquitin chains on target proteins are primarily associated with proteasomal degradation, whereas K63-linked mono- or polyubiquitin chains do not promote degradation but instead alter the biochemical properties of proteins, leading to different cellular outcomes, we aimed to distinguish which type of ubiquitination is involved in this process. To identify the type of ubiquitin linkage, cells were transfected with ubiquitin mutants allowing only K48- or K63-linked chains. Co-IP revealed that DAPK1 specifically promoted K48-linked, but not K63-linked, polyubiquitination of parkin (Fig. 2D), consistent with proteasomal targeting.

Collectively, these results demonstrate that DAPK1 facilitates parkin degradation by promoting its K48-linked polyubiquitination and subsequent proteasomal degradation via a kinase-dependent mechanism.

### DAPK1 phosphorylates parkin on S136 and S198 residues

Given that DAPK1-mediated parkin degradation requires its kinase activity, we investigated whether DAPK1 directly phosphorylates parkin. Phos-tag immunoblot analysis revealed parkin phosphorylation in the presence of DAPK1-WT, but not with the kinase-defective DAPK1-KD, indicating a kinase-dependent modification (Fig. 3A). An *in vitro* kinase assay using purified recombinant proteins confirmed that parkin is phosphorylated by DAPK1-WT, but not DAPK1-KD, supporting a direct phosphorylation event (Fig. 3B).

**Fig. 3.**
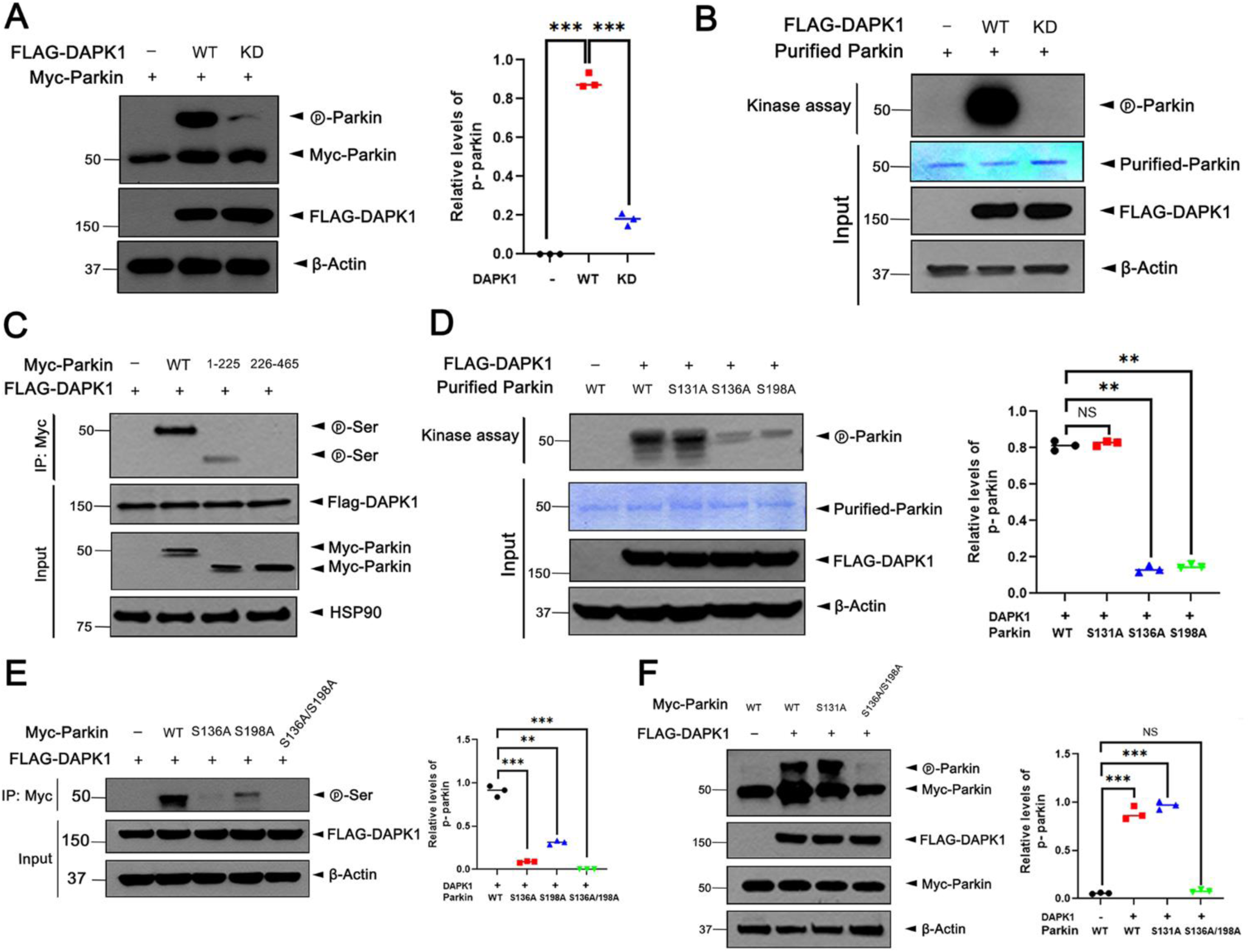
DAPK1 phosphorylates parkin on S136 and S198 residues. (A) MN9D cells were transfected with Myc-parkin for 24 h individually or in combination with FLAG-DAPK1-WT or FLAG-DAPK1-KD. Cell lysates were resolved by Phos-tag gel electrophoresis and immunoblotted with the indicated antibodies. Quantification of relative phosphorylated parkin expression was quantified (mean ± SD, n = 3; (\*\*\**p* ≤ 0.0001). (B) FLAG-DAPK1-WT or -KD was immunoprecipitated from MN9D cells and incubated in kinase buffer with recombinant GST-cleaved parkin and [γ-^32^P] for 2 h at 37°C. The reaction mixtures were analyzed by SDS-PAGE followed by autoradiography. Coomassie brilliant blue (CBB) staining confirmed protein inputs. (C) MN9D cells were transfected with FLAG-DAPK1 alone or co-transfected with Myc-parkin-WT, Myc-parkin-1–225, or Myc-parkin-226–465 for 24 h. Cell lysates were examined by immunoblotting with the indicated antibodies. (D) FLAG-DAPK1-WT immunocomplexes were incubated with recombinant GST-parkin or its point mutants (S131A, S136A, or S198A) in kinase buffer with [γ-^32^P]. Reaction products were subjected to SDS-PAGE and analyzed by autoradiography. Protein loading was confirmed by CBB staining. Quantification of relative phosphorylated parkin expression was quantified (mean ± SD, n = 3; \*\**p* ≤ 0.001; N.S., not significant). (E, F) MN9D cells expressing FLAG-DAPK1 together with Myc-parkin-WT, Myc-parkin-S131A, Myc-parkin-S136A, Myc-parkin-S198A, or the double mutant (S136A/S198A) were analyzed by immunoblotting (E) or Phos-tag gels (F) using the indicated antibodies. Quantification of relative phosphorylated parkin expression was quantified (mean ± SD, n = 3; E: \*\*\**p* ≤ 0.0001, \*\**p* ≤ 0.001; F: \*\*\**p* ≤ 0.0001, N.S., not significant). Hsp90 and β-actin served as internal controls for protein loading.

To map the phosphorylation site(s), two parkin truncation mutants (1–225 and 226–465) were co-expressed with DAPK1-WT in MN9D cells. Phospho-serine immunoblotting showed phosphorylation in parkin-WT and the N-terminal 1–225 mutant, but not in the C-terminal 226–465 fragment, indicating the site resides within the N-terminal region (Fig. 3C).

To further narrow down the precise phosphorylation sites within the parkin-1-225 region, three well-characterized serine residues on parkin were selected for mapping. These included two serine residues (S131 and S136), known targets of other kinases in parkin, as well as S198, a potential phosphorylation site inferred from analysis of DAPK1 substrates. *In vitro* kinase assays revealed that S136A and S198A mutants displayed significantly reduced phosphorylation, while S131A remained unaffected (Fig. 3D). Notably, the double mutant S136A/S198A exhibited a complete loss of phosphorylation, while S136A alone showed minimal phosphorylation compared to the relatively higher levels in S198A (Fig. 3E). These findings suggest that S136 is the primary phosphorylation site, with S198 as a secondary site.

Finally, Phos-tag analysis confirmed robust phosphorylation in parkin-WT and S131A, but not in the S136A/S198A mutant (Fig. 3F), reinforcing that S136 and S198 are critical DAPK1-targeted residues. Collectively, these results demonstrate that DAPK1 directly phosphorylates parkin at S136 and S198, providing mechanistic insight into how DAPK1 regulates parkin stability and function.

### DAPK1-mediated phosphorylation of parkin at S136 and S198 enhances its degradation

To determine whether DAPK1-mediated parkin degradation is driven by phosphorylation at S136/S198, we used a double phosphorylation-resistant mutant (parkin-S136A/S198A; parkin-2A). In MN9D cells, co-expression of DAPK1-WT significantly reduced wild-type parkin levels, but not parkin-2A levels, indicating that phosphorylation at these residues is essential for DAPK1-induced degradation (Fig. 4A). Similarly, DAPK1 enhanced polyubiquitination of parkin-WT but not parkin-2A, confirming the requirement of phosphorylation for ubiquitination (Fig. 4B). A phosphorylation-mimetic mutant (parkin-S136E/S198E; parkin-2E) exhibited increased polyubiquitination relative to wild type, further supporting that phosphorylation at these sites promotes ubiquitination and degradation (Fig. 4C).

**Fig. 4.**
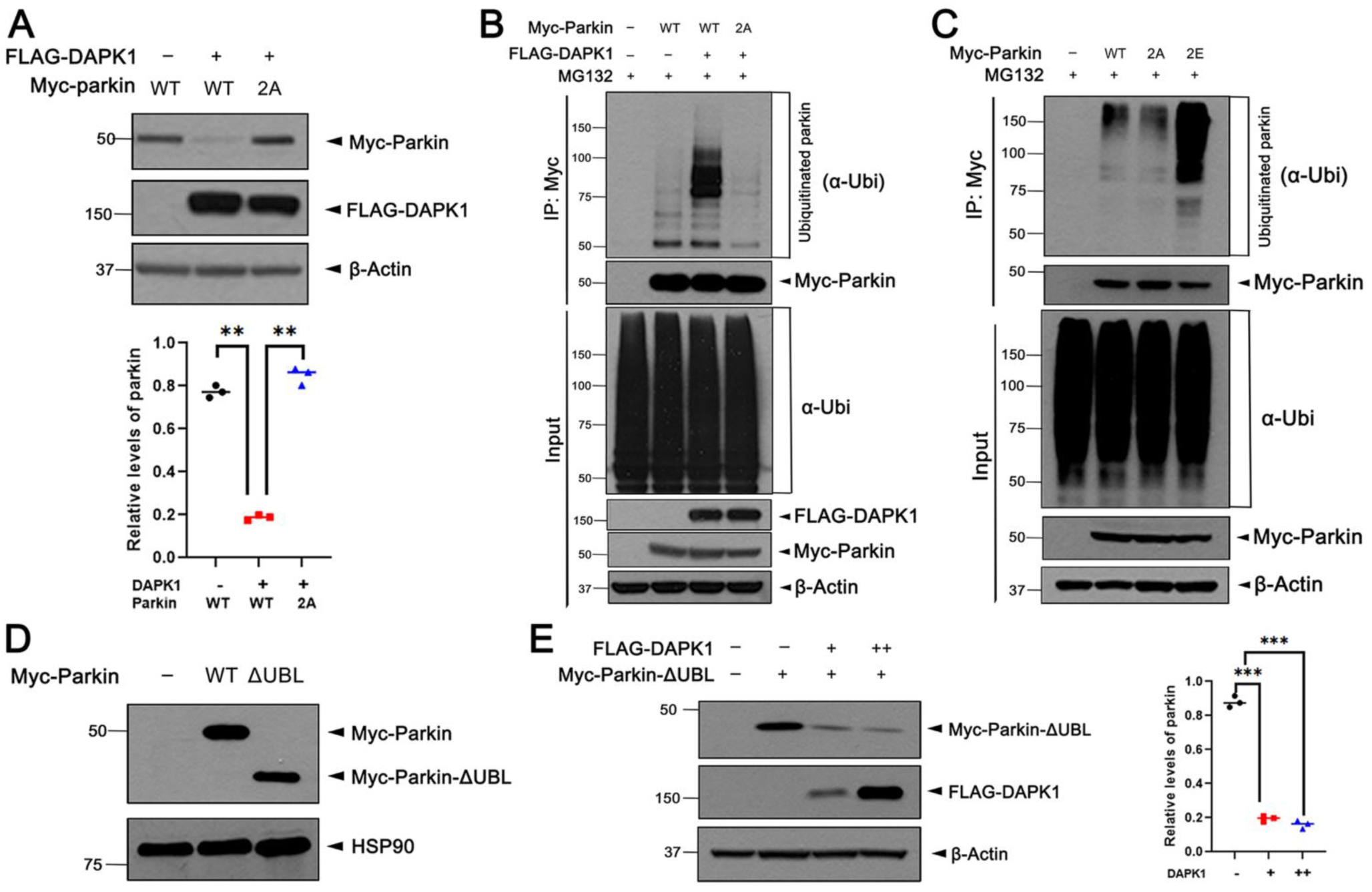
DAPK1-mediated phosphorylation of parkin at S136 and S198 enhances its degradation. (A) MN9D cells were transfected for 24 h with FLAG-DAPK1, Myc-parkin-WT, or the phospho-deficient mutant Myc-parkin-S136A/S198A (parkin-2A), either alone or in combination. Lysates were immunoblotted, and parkin expression was quantified (mean ± SD, n = 3; \*\**p* ≤ 0.001). (B) Cells expressing FLAG-DAPK1 and either parkin-WT or parkin-2A were treated with 10 μM MG132 for 6 h. Myc-immunoprecipitates were analyzed by immunoblotting using the indicated antibodies. (C) To assess phosphorylation-mimicking effects, MN9D cells expressing parkin-WT, parkin-2A, or parkin-2E were treated with MG132. Cell lysates were immunoprecipitated with anti-Myc antibody, and Myc-immunoprecipitates were analyzed by immunoblotting using the indicated antibodies. (D) To examine the contribution of the UBL domain, MN9D cells were transfected with Myc-parkin-WT or Myc-parkin-ΔUBL, and lysates were immunoblotted with the indicated antibodies. (E) Increasing amounts of FLAG-DAPK1 were co-expressed with Myc-parkin-ΔUBL for 24 h. Cell lysates were immunoblotted with the indicated antibodies. Quantification of relative parkin- ΔUBL expression was quantified (mean ± SD, n = 3; \*\*\**p* ≤ 0.0001). Hsp90 and β-actin served as internal controls for protein loading.

To assess whether this degradation depends on parkin’s own E3 ligase activity, we examined a dominant-negative mutant lacking the UBL domain (parkin-ΔUBL), essential for its ubiquitination function (Chaugule et al., 2011). DAPK1 still reduced parkin-ΔUBL levels, indicating that degradation is independent of parkin’s auto-ubiquitination and instead driven by DAPK1-mediated phosphorylation (Fig. 4D, E).

Together, these findings demonstrate that DAPK1 phosphorylates parkin at S136/S198, facilitating its polyubiquitination and proteasomal degradation via a mechanism likely involving other UPS components.

### Treatment of MN9D cells with 6-OHDA causes the reduction of parkin via DAPK1-mediated phosphorylation

Given that DAPK1-mediated phosphorylation leads to parkin degradation, we hypothesized that DAPK1 may impair parkin’s neuroprotective role under oxidative stress, contributing to neuronal vulnerability in PD (Cheung & IP, 2012). To test this, MN9D cells were treated with various PD-related neurotoxins. Among them, only 6-OHDA significantly reduced parkin protein levels, suggesting selective sensitivity (Fig. S3A). DAPK1 overexpression further enhanced 6-OHDA-induced parkin reduction, indicating a synergistic effect (Fig. S3B). This effect was further examined using parkin-2A and parkin-2E mutants. Notably, parkin-2A was resistant to 6-OHDA-induced degradation, while parkin-2E showed enhanced reduction (Fig. S3C), confirming the critical role of S136/S198 phosphorylation in this process.

Endogenous parkin levels were also decreased by 6-OHDA treatment (Fig. S3D). However, *DAPK1*-knockdown using siRNA partially rescued this reduction, demonstrating that DAPK1 is necessary for 6-OHDA-induced parkin degradation (Fig. S3E). Since 6-OHDA induces dopaminergic cell death through oxidative stress and is widely used in PD models (Moore et al., 2005), these findings suggest that DAPK1 contributes to PD pathology by facilitating parkin degradation under oxidative stress. Thus, DAPK1 may act as a crucial mediator linking ROS-induced damage to parkin inactivation in dopaminergic neurons.

### DAPK1-mediated phosphorylation of parkin promotes its mitochondrial transport and enhances the binding between parkin and ubiquitin E3 ligase MITOL

To identify the UPS component responsible for DAPK1-mediated parkin degradation, we focused on MITOL (also known as MARCH5), a mitochondrial E3 ligase previously reported to interact with parkin under mitophagic conditions (Shiiba et al., 2021). We hypothesized that DAPK1-induced phosphorylation may facilitate MITOL-dependent parkin degradation. Co-IP analysis revealed that under basal conditions, parkin and MITOL did not interact; however, DAPK1 overexpression significantly enhanced their binding (Fig. 5A). To examine whether this interaction depends on parkin phosphorylation, we compared MITOL binding to wild-type parkin, the phospho-mimetic parkin-2E, and the phosphorylation-resistant parkin-2A mutant. Notably, MITOL interacted strongly with parkin-2E but not with parkin-2A or wild-type parkin, indicating that DAPK1-mediated phosphorylation enhances parkin-MITOL association (Fig. 5B).

**Fig. 5.**
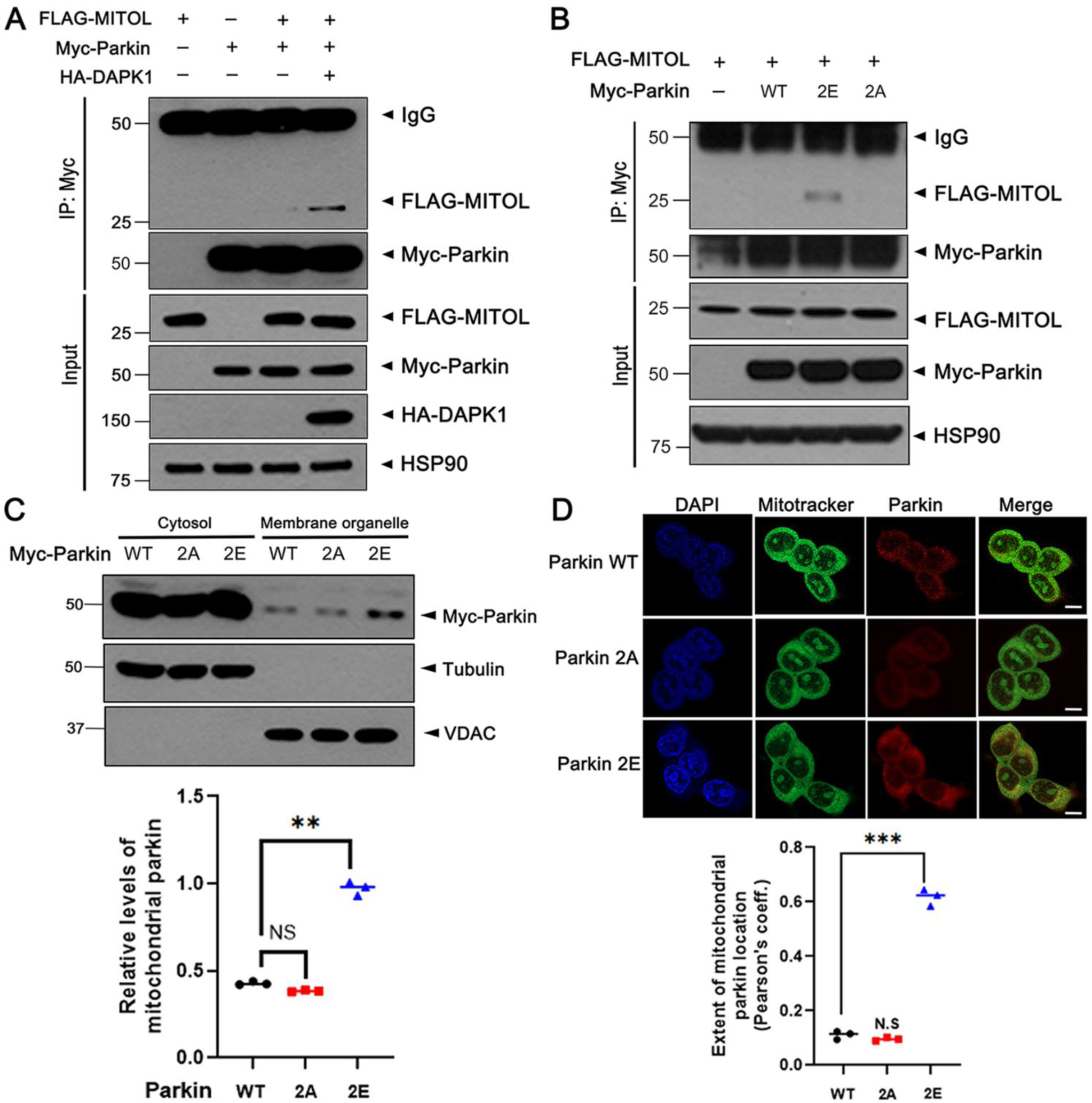
DAPK1-mediated proteolysis of parkin occurs through mitochondrial transport of parkin and subsequent action of MITOL under 6-OHDA treatment in MN9D cells. (A) MN9D cells were transfected for 24 h with Myc-parkin, HA-DAPK1, or FLAG-MITOL, alone or in combination, and treated with 10 μM MG132 for 6 h. Myc-immunoprecipitates were analyzed by immunoblotting with the indicated antibodies. (B) FLAG-MITOL was co-expressed with Myc-parkin-WT, parkin-2A, or parkin-2E. After treatment with MG132 for 6 h, lysates were immunoprecipitated using anti-Myc antibody, and the immunoprecipitates were analyzed by immunoblotting with the indicated antibodies. (C) MN9D cells expressing Myc-parkin-WT or its mutants (2A, 2E) were fractionated into cytosolic and membrane fractions. Each fraction was immunoblotted with the indicated antibodies. Quantification of relative mitochondrial parkin enrichment expression was quantified (mean ± SD, n = 3; \*\**p* ≤ 0.001; NS., not significant). (D) MN9D cells were subjected to immunofluorescence staining. Confocal images demonstrate colocalization of parkin (red) with MitoTracker (green), while nuclei were stained with DAPI (blue). Scale bars, 10 μm. Quantification of relative mitochondrial parkin enrichment expression was quantified (mean ± SD, n = 3; \*\*\**p* ≤ 0.0001; NS., not significant). Hsp90, VDAC, and tubulin were used as fractionation/loading controls.

Given MITOL’s mitochondrial localization, we tested whether phosphorylation promotes parkin translocation to mitochondria. Subcellular fractionation experiments demonstrated that parkin-2E showed a 2.2-fold increase in mitochondrial localization compared to parkin-WT and parkin-2A (Fig. 5C). This observation was further validated using immunostaining, where parkin localization was analyzed using an antibody against parkin and MitoTracker, a selective mitochondrial dye; notably, parkin-2E displayed more than a sixfold increase in colocalization with mitochondria than the other variants (Fig. 5D).

Collectively, these findings demonstrate that DAPK1-catalyzed phosphorylation of parkin facilitates its translocation to mitochondria, where it interacts with MITOL, leading to proteasomal degradation. This identifies MITOL as a likely mediator of DAPK1-dependent parkin turnover and highlights a potential regulatory mechanism linking kinase activity, subcellular localization, and protein stability.

### The cellular toxicity of 6-OHDA in MN9D cells is attributed to DAPK1-mediated phosphorylation and proteolysis of parkin

Parkin is a well-established neuroprotective protein that defends dopaminergic neurons against a variety of toxic insults, including MPTP, 6-OHDA, and rotenone (Shiiba et al., 2021). To assess whether phosphorylation by DAPK1 diminishes parkin’s protective capacity, we investigated its involvement in 6-OHDA-induced cytotoxicity using MN9D cells. Treatment with 6-OHDA progressively reduced cell viability, and exposure to 100 μM resulted in nearly 50% cell loss, as determined by CCK-8 analysis (Fig. 6A). Overexpression of Myc-parkin-WT significantly rescued cell viability, whereas co-expression of FLAG-DAPK1-WT reversed this protective effect. In contrast, the kinase-dead mutant DAPK1-KD did not affect viability, suggesting that DAPK1’s kinase activity is essential for suppressing parkin-mediated protection (Fig. 6A). Consistently, knockdown of DAPK1 enhanced parkin-dependent cell survival under 6-OHDA stress (Fig. 6B).

**Fig. 6.**
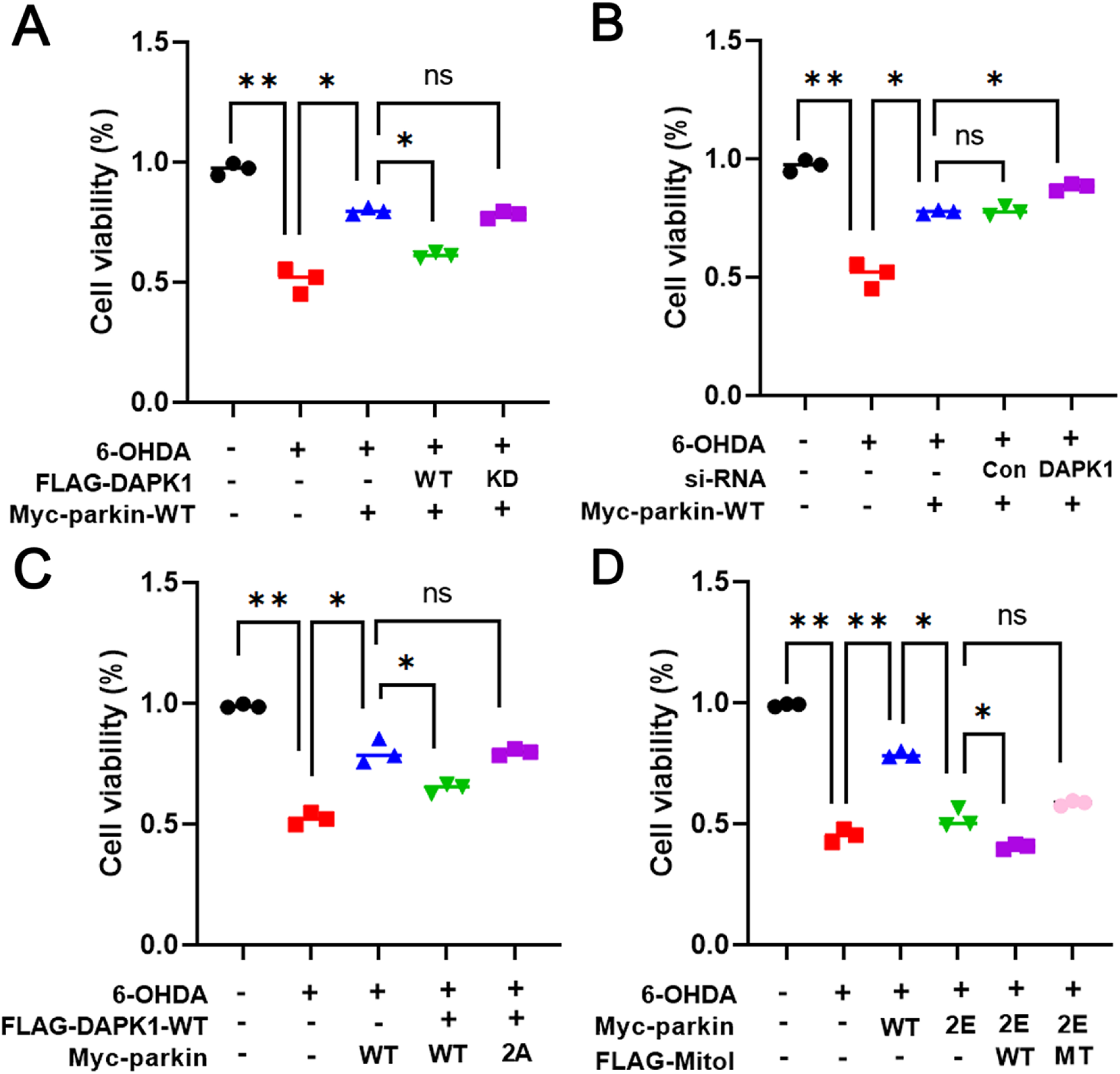
DAPK1-mediated phosphorylation inhibits the neuroprotective effect of parkin against 6-OHDA in MN9D cells. (A) MN9D cells were transfected for 24 h with Myc-parkin, FLAG-DAPK1-WT, or FLAG-DAPK1-KD, either individually or in combination. Cells were then treated with 100 μM 6-OHDA for 24 h, and viability was determined by CCK-8 assay. Quantification of relative parkin level was quantified (mean ± SD, n = 3; **p ≤ 0.001; \**p* ≤ 0.05; NS., not significant). (B) MN9D cells were transfected with Myc-parkin, scrambled siRNA (control), or *DAPK1*-specific siRNA, and then treated with 100 μM 6-OHDA for 24 h. Cell viability was measured using CCK-8 assays (mean ± SD, n = 3; \*\**p* ≤ 0.001; \**p* ≤ 0.05; NS., not significant). (C) Where indicated, MN9D cells were transfected for 24 h with Myc-parkin-WT, Myc-parkin-2A, or FLAG-DAPK1-WT. Cells were treated with 100 μM 6-OHDA for 24 h, and viability was assessed by CCK-8 assays (mean ± SD, n = 3; \*\**p* ≤ 0.001; *p ≤ 0.05; NS., not significant). (D) MN9D cells were transfected with Myc-parkin-WT, Myc-parkin-2E, FLAG-MITOL-WT, or FLAG-MITOL-MT for 24 h, either alone or in combination. Cells were then treated with 100 μM 6-OHDA for 24 h, and viability was measured using CCK-8 assay. (mean ± SD, n = 3; \*\**p* ≤ 0.001; \**p* ≤ 0.05; NS., not significant).

To determine whether parkin phosphorylation is required for this effect, we compared the impact of DAPK1 on cells expressing either parkin-WT or parkin-2A mutant. Notably, the reduced neuroprotective effect of parkin observed upon DAPK1 overexpression was restored in cells expressing parkin-2A, supporting a phosphorylation-dependent mechanism (Fig. 6C).

Finally, we evaluated the role of MITOL in this process. Overexpression of the phospho-mimetic parkin-2E markedly increased susceptibility to 6-OHDA-induced toxicity, indicating a loss of its neuroprotective function. This effect was further exacerbated by co-expression of MITOL-WT, consistent with its role in promoting parkin degradation. In contrast, the catalytically inactive MITOL mutant (MITOL-MT), which carries C65S and C68S substitutions that abolish its E3 ligase activity, failed to enhance 6-OHDA-induced toxicity, highlighting the requirement of MITOL’s enzymatic function for mediating parkin degradation and the resulting neurotoxicity (Fig. 6D). These results suggest that DAPK1-mediated phosphorylation of parkin promotes its MITOL-dependent degradation, leading to loss of neuroprotection and increased vulnerability to neurotoxic stress.

Together, these findings reveal a novel mechanism by which DAPK1 impairs parkin’s protective role, contributing to dopaminergic neurodegeneration in PD models.

## Discussion

Phosphorylation, a key post-translational modification, regulates parkin by modulating its activity and stability via various kinases, including Dyrk1A, PINK1, c-Abl, and CK2. For instance, PINK1-mediated phosphorylation at Ser65 activates parkin, facilitating its mitochondrial recruitment and mitophagy initiation (Narendra et al., 2010). This modification enhances parkin’s E3 ligase activity, promoting ubiquitination of outer mitochondrial membrane proteins and the clearance of damaged mitochondria (Pickrell & Youle, 2015). Conversely, phosphorylation at Tyr143 by c-Abl suppresses parkin’s ligase activity, leading to mitochondrial dysfunction and increased neurotoxicity in PD models (Zhang et al., 2022). In this context, our study newly identifies DAPK1 as a kinase that phosphorylates parkin, not to activate it, but to promote its degradation, thereby regulating its protein stability under neurotoxic conditions. These findings expand the understanding of parkin regulation and highlight the diverse consequences of phosphorylation. The intricate network of parkin PTMs reflects its physiological importance, as dysregulation of parkin activity is strongly linked to both PD pathogenesis and tumorigenesis.

DAPK1 has multiple substrates that mediate distinct physiological outcomes. For example, DAPK1 phosphorylates p53, stabilizing it by preventing MDM2-mediated degradation. This leads to upregulation of pro-apoptotic genes like Bax and PUMA, promoting cytochrome c release and caspase activation, which contribute to neuronal loss in NDDs (Cheung & Ip, 2012; Moore et al., 2005). DAPK1 also targets Beclin-1, a key autophagy regulator; its phosphorylation disrupts Beclin-1’s interaction with Bcl-2, enhancing autophagy and promoting neuronal survival under stress (Tilija & Park, 2018). Given the link between impaired autophagy and PD, these findings suggest that DAPK1 modulates neuronal homeostasis via autophagy pathways. Furthermore, we previously reported that DAPK1 phosphorylates α-synuclein, increasing its aggregation and toxicity (Shin & Chung, 2020). In addition to α-synuclein, our current study identifies parkin as another critical DAPK1 target. DAPK1-mediated phosphorylation of parkin promotes its degradation through the UPS, revealing a novel mechanism by which DAPK1 contributes to PD pathogenesis.

Given parkin’s critical role in mitophagy and neuroprotection, its protein stability is tightly regulated. Parkin can undergo auto-ubiquitination and is also degraded by UPS components such as MITOL, a mitochondrial E3 ligase (Chaugule et al., 2011). MITOL-mediated proteolysis of parkin is heightened under stress conditions like oxidative damage, suggesting MITOL acts as a sensor to modulate parkin levels. Although parkin’s degradation via other E3 ligases or the autophagy-lysosome pathway remains poorly understood, it is known that parkin-mediated K63-linked polyubiquitination facilitates the sequestration of misfolded proteins into aggresomes for autophagic clearance (Olzmann & Chin, 2008). In this study, we reveal a novel mechanism whereby 6-OHDA treatment and subsequent DAPK1-mediated phosphorylation induce parkin’s translocation to mitochondria. This enhances its interaction with MITOL, resulting in its degradation.

Similar to DAPK1-mediated parkin transport, phosphorylation is known to regulate the mitochondrial translocation of various cytosolic proteins. For example, the pro-apoptotic protein BAD translocates to mitochondria upon dephosphorylation, as 14-3-3 proteins normally sequester it in the cytosol (Datta et al., 2000). Once dephosphorylated, BAD dissociates from 14-3-3 and binds Bcl-2 family proteins to trigger apoptosis (Zha et al., 1996). Conversely, phosphorylation by survival kinases like Akt and PKA retains BAD in the cytosol (del Peso et al., 1997). Similarly, p53, usually localized in the nucleus and cytosol, translocates to mitochondria following phosphorylation by HIPK2 or ATM/ATR in response to DNA damage, where it promotes apoptosis by interacting with Bcl-2 family proteins (Mihara et al., 2003). Based on these precedents, DAPK1-induced phosphorylation of parkin may expose a mitochondrial targeting motif or alter regulatory interactions, facilitating its mitochondrial localization and degradation. Further studies are needed to define the precise structural and interaction changes driving this process.

Our results show that DAPK1-mediated phosphorylation promotes parkin degradation through a MITOL-dependent mechanism, linking phosphorylation to mitochondrial quality control. Phosphorylation at S136 and S198 enhances MITOL binding, with S136 having a dominant effect. Under basal conditions, MITOL does not interact with non-phosphorylated parkin, but binding is significantly increased by DAPK1 overexpression or 6-OHDA treatment. Supporting this, MITOL strongly binds only to the phospho-mimetic 2E mutant, not to the phosphorylation-resistant 2A variant, indicating that parkin phosphorylation is essential for its recognition and degradation by MITOL.

To understand the structural basis of this phenomenon, we performed protein structure prediction analysis. The results suggest that S136 phosphorylation alters parkin’s conformation, exposing a positively charged patch in the RING0 domain (K161, R163, R455), which may facilitate MITOL binding (Fig. S4A and B). Additionally, S198 phosphorylation likely disrupts its hydrogen bonding with Q171 in the RING0 domain, potentially destabilizing parkin’s structure and enhancing its degradation (Fig. S4C and D). Given the key role of the RING0 domain in parkin activation, these findings suggest that DAPK1-induced phosphorylation might impair parkin’s E3 ligase activity, further compromising its ability to exert neuroprotective effects. It would be interesting to experimentally test whether mutating these positively charged residues (K161, R163, R455) to negatively charged residues (D or E) affects the impact of S136 phosphorylation, as this could provide additional insight into the structural basis of parkin’s degradation. Furthermore, AlphaFold3-based structural modeling of the Ser136-phosphorylated parkin complex suggests that phosphoserine 136 interacts with key basic residues, providing a molecular explanation for how phosphorylation alters parkin’s structure and function. Notably, these interactions may facilitate the recruitment of MITOL and other degradation-associated factors. Moreover, the effect of S136 phosphorylation is expected to be reduced if key positively charged residues (K161, R163, R455) are mutated to negatively charged amino acids, supporting the hypothesis that charge-based interactions contribute to parkin’s stability and degradation.

6-OHDA, a well-established dopaminergic neurotoxin, is widely used in PD models to study neuronal degeneration. Upon exposure, it elevates ROS levels, induces mitochondrial dysfunction, and activates intrinsic apoptotic pathways (Liang et al., 2004). It also promotes α-synuclein aggregation, leading to intracellular inclusions in dopaminergic neurons (Zhang et al., 2022). Based on these findings and our data, we propose that 6-OHDA-induced neurotoxicity is mediated by toxic protein aggregates. This is supported by our observation that DAPK1 overexpression aggravates, whereas its knockout alleviates, 6-OHDA-induced neuronal loss. Furthermore, we identified DAPK1 as a dual contributor to PD pathogenesis: it phosphorylates α-synuclein at Ser129 to enhance aggregation (Shin & Chung, 2020), and also promotes parkin degradation. Since parkin is essential for mitophagy and clearance of misfolded proteins, its loss may impair mitochondrial quality control and exacerbate α-synuclein pathology. Thus, DAPK1-induced parkin degradation could further promote protein aggregation and neurodegeneration, contributing to PD progression.

Overall, this study provides compelling evidence that DAPK1-mediated phosphorylation serves as a key regulatory event in parkin degradation, linking it to neurotoxicity in PD. By identifying phosphorylation as an alternative mechanism of parkin degradation, we offer a new perspective on PD pathogenesis and potential therapeutic strategies for preserving parkin function.

## Data availability

The datasets generated and/or analyzed during this work are available from the corresponding author when requested reasonably.

## Acknowledgements

We are grateful to T.H. Lee, K. Tanaka, and S. Hirose for generously sharing plasmids, and to T.H. Lee for additionally providing MEFs derived from DAPK1-deficient mice.

## Author contributions

Chul Hong Park performed DNA transfection, *in vitro* GST pull-down, *in vitro* kinase assays, and CCK assays, western blotting, and data analysis. He was also responsible for drafting the manuscript. Donghyuk Shin contributed by performing AlphaFold structural predictions and assisting with data interpretation. Kwang Chul Chung conceived and designed the study, supervised experimental execution, interpreted the results, wrote parts of the manuscript, secured funding, and took full responsibility for the study.

## Supplementary material

This article contains Supplementary material.

