## Supplementary Data for "DAPK1-mediated parkin inactivation enhances neurotoxicity via MITOL-dependent degradation"

### Supplementary Figures

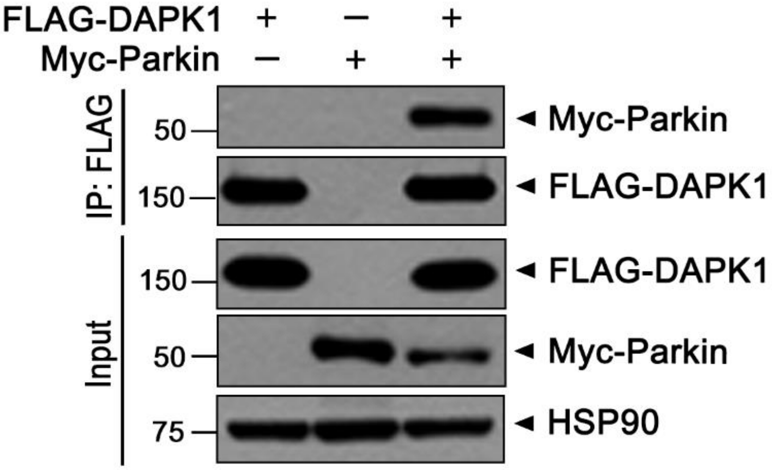


**Fig. S1 Co-immunoprecipitation reveals interaction between parkin and DAPK1 in the absence of MG132.** Where indicated, HEK293 cells were transfected for 24 h with plasmids encoding Myc-parkin or FLAG-DAPK1 alone or in combination. Cell lysates were subjected to immunoprecipitation using an anti-FLAG antibody, and the resulting immunoprecipitates were immunoblotted with the indicated antibodies.

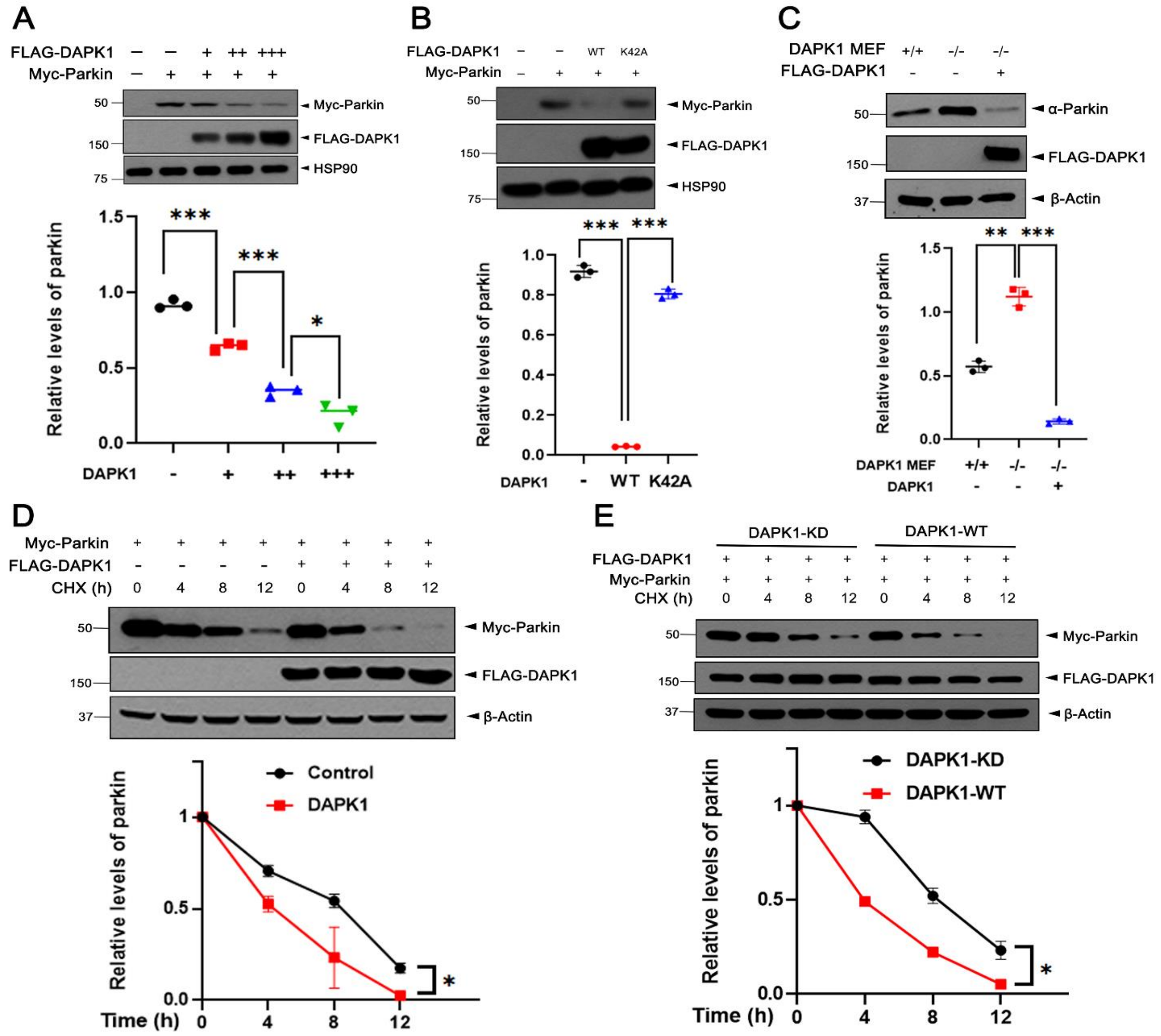


**Fig. S2 DAPK1 decreases the protein stability of parkin.** (A) MN9D cells were transfected for 24 h with plasmids encoding Myc-parkin alone or together with increasing amounts of FLAG-DAPK1. Cell lysates were immunoblotted with the indicated antibodies. Relative levels of parkin were quantified and the results presented as the mean ± SD of three independent experiments (***$p \leq 0.0001$; **$p \leq 0.001$). (B) MN9D cells were transfected for 24 h with plasmids encoding Myc-parkin alone or in combination with FLAG-DAPK1-WT or FLAG-DAPK1-K42A. Cell lysates were immunoblotted with the indicated antibodies. Relative levels of parkin were quantified and results presented as the mean ± SD of three independent experiments (***$p \leq 0.0001$). (C) The mouse embryo fibroblasts (MEF) expressing wild-type DAPK1 (DAPK1-WT) or its knockout (*DAPK1*-KO) MEFs were mock-transfected or transfected for 24 h with

FLAG DAPK1. Cell lysates were immunoblotted with the indicated antibodies. Relative levels of parkin were quantified and the results presented as the mean ± SD of three independent experiments (***$p \leq 0.0001$; **$p \leq 0.001$). (D) MN9D cells were transfected for 24 h with plasmids encoding Myc-parkin alone or together with FLAG-DAPK1. Cells were treated for the indicated times with 25 μg/ml cycloheximide (CHX), and cell lysates were immunoblotted with the indicated antibodies. Relative levels of parkin were quantified and the results presented as the mean ± SD of three independent experiments (*$p \leq 0.05$). (E) Where indicated, MN9D cells were transfected for 24 h with plasmids encoding Myc-parkin, along with either FLAG-DAPK1-KD or FLAG-DAPK1-WT. Cells were treated for the indicated times with 25 μg/ml CHX, and cell lysates were immunoblotted with the indicated antibodies. Relative levels of parkin were quantified and the results presented as the mean ± SD of three independent experiments (*$p \leq 0.05$). β-Actin and Hsp90 served as a loading control.

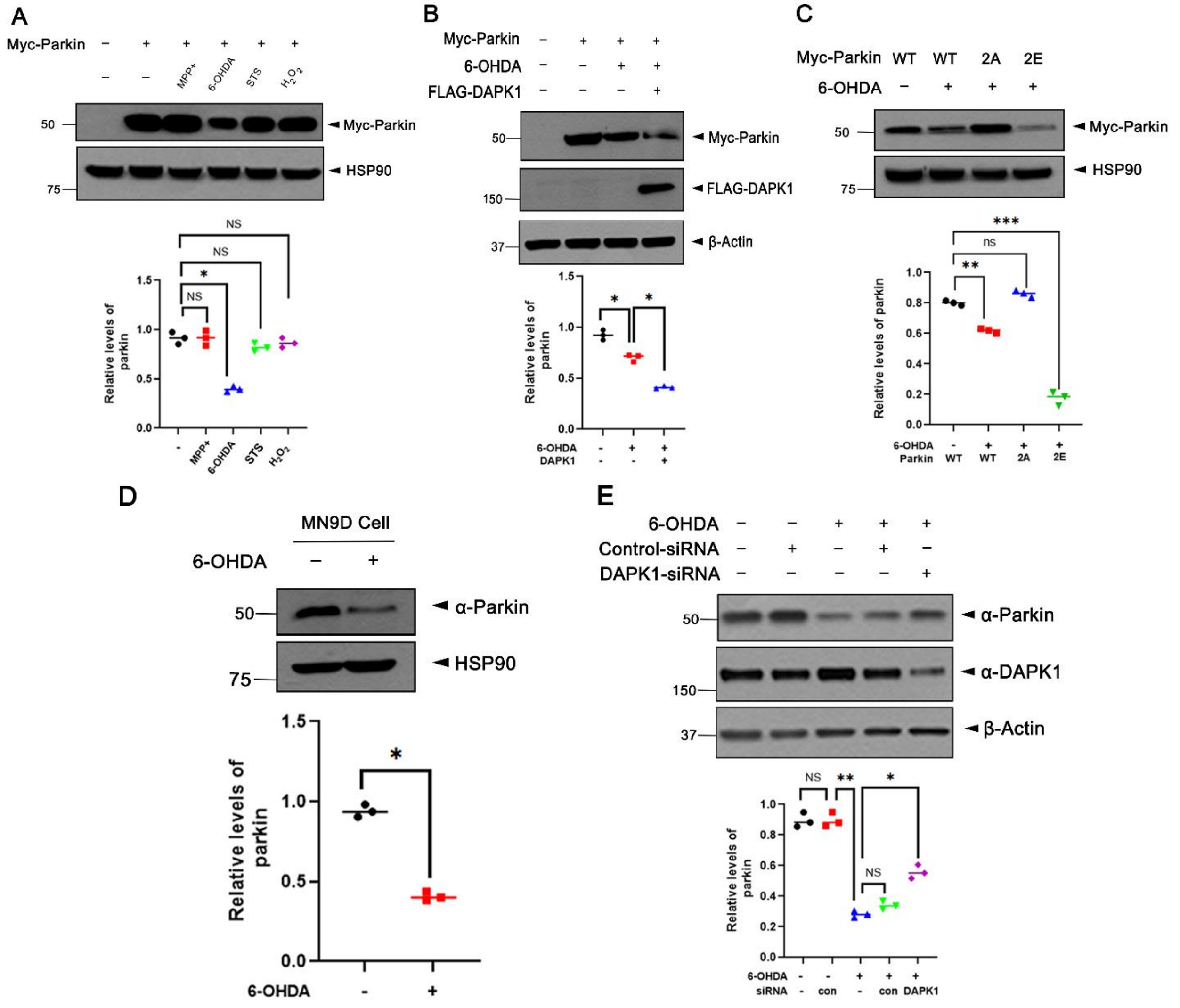


**Fig. S3 Exposure of MN9D cells to 6-OHDA caused the reduction of parkin.** (A) Where indicated, MN9D cells were treated for 6 h with $MPP^+$ (500 μM), 6-OHDA (50 μM), staurosporine (STS, 0.5 μM), or $H_2O_2$ (500 μM). Cell lysates were immunoblotted with the indicated antibodies. Relative parkin levels were quantified and are presented as the mean ± SD of three independent experiments (*$p \leq 0.05$; NS, not significant). (B) MN9D cells were transfected for 24 h with plasmids encoding FLAG-DAPK1 alone or in combination with Myc-parkin, and treated for additional 6 h with 50 μM 6-OHDA. Cell lysates were immunoblotted with the indicated antibodies. Relative parkin levels were quantified and are presented as the mean ± SD of three independent experiments (*$p \leq 0.05$). (C) MN9D cells were transfected for 24 h with Myc-parkin-WT, Myc-parkin-2A, or Myc-parkin-2E and treated for additional 6 h with 50 μM 6-OHDA. Cell lysates were immunoblotted with the indicated antibodies. Relative parkin levels were quantified and are presented as the mean ± SD of three independent experiments (***$p \leq 0.0001$; **$p \leq$

0.001; NS, not significant). (D) MN9D cells were treated for 6 h with vehicle (-) or 50 μM 6-OHDA. Cell lysates were immunoblotted with the indicated antibodies. Relative parkin levels were quantified and are presented as the mean ± SD of three independent experiments (*$p \leq 0.05$). (E) MN9D cells were transfected for 48 h with control siRNA or *DAPK1*-siRNA. Cells were then left untreated or treated for additional 4 h with 50 μM 6-OHDA, and cell lysates were immunoblotted with the indicated antibodies. Relative parkin levels were quantified and are presented as the mean ± SD of three independent experiments (**$p \leq 0.001$; *$p \leq 0.05$; NS, not significant). β-Actin and Hsp90 served as a loading control.

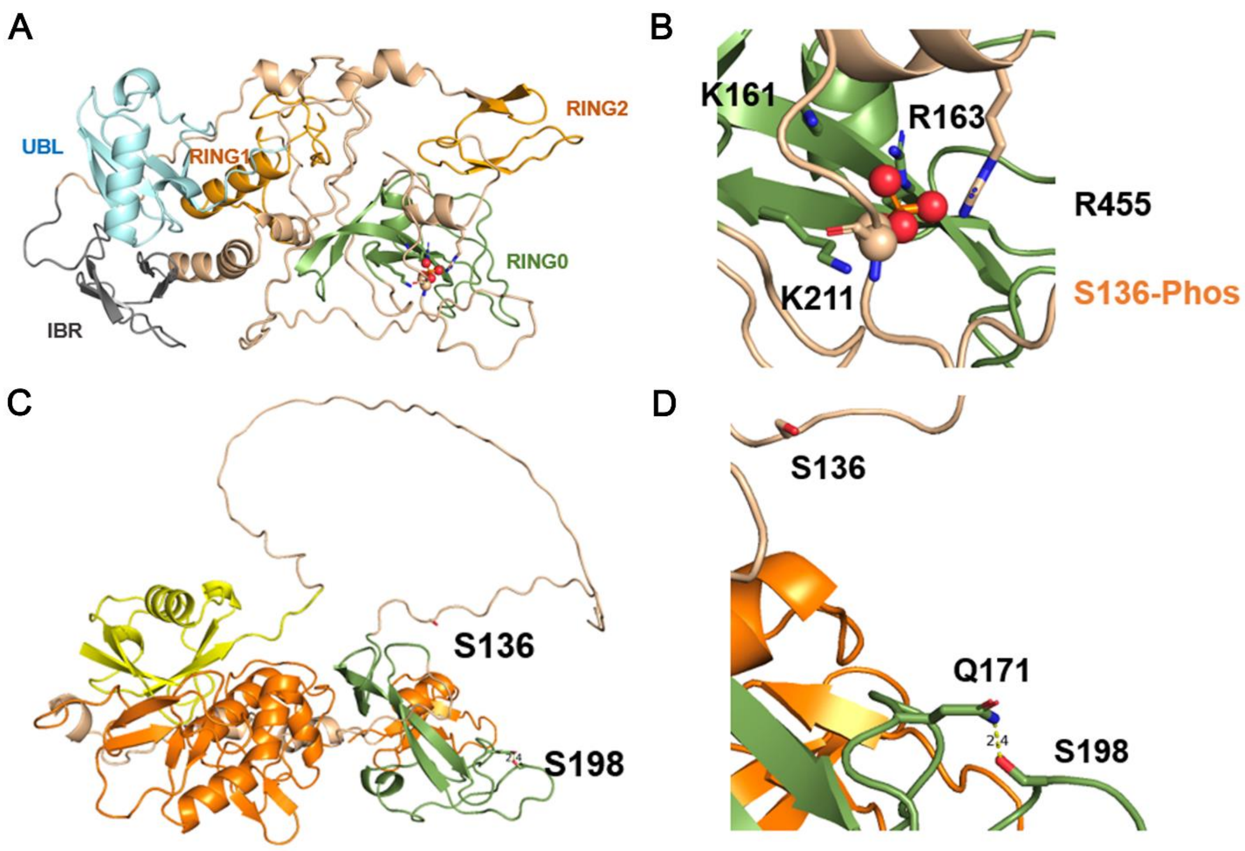


**Fig. S4 Predicted structure of phosphorylated Parkin.** (A, B) Ser136 and Ser198 of Parkin are highlighted in the predicted Parkin structure. Ser136 is located on the loop between UBL and RING0, while Ser198 forms a hydrogen bond with Q171. (C) Representative model of the AlphaFold3-predicted complex structure of Ser136-phosphorylated Parkin. (D) Basic amino acids interacting with phosphoserine 136 are highlighted.
